# Curved by Design: Applying Microfluidic Principles for Nonplanar and Planar Suspended Tissue Patterning to the Development of Bladder Models with Tunable Mechanics

**DOI:** 10.64898/2026.09.11.751043

**Authors:** Jamison M. Whitten, Ariel Lin, Ella E. Bouker, Asha R. Viswanathan, Jean Berthier, Xiaojing Su, Emme A. Schumacher, Erwin Berthier, Ashleigh B. Theberge, Amanda J. Haack, Maya R. Chandru

**Affiliations:** Department of Chemistry, University of Washington, Seattle, WA, 98195 USA; Molecular Engineering & Sciences Institute, University of Washington, Seattle, WA, 98109 USA; Department of Urology, University of Washington School of Medicine, Seattle, WA, 98195 USA; Department of Chemical Engineering, Texas Tech University, Lubbock, TX, 79409 USA; Department of Bioengineering, University of Washington, Seattle, WA, 98195 USA; Division of Pediatric Urology, Seattle Children’s Hospital, Seattle, WA, 98105 USA

## Abstract

Cells *in vivo* exist in a complex environment where they receive chemical and physical cues from neighboring cells and the extracellular matrix. Suspended, three dimensional (3D) cell culture enables the study of mechanical signals in a controlled *in vitro* setting where cells can exert forces on the extracellular matrix, and mechanical stimulation can be externally applied. In addition to mechanical cues, tissues *in vivo* also exhibit spatial heterogeneity and nonplanar topography. To facilitate the development of suspended 3D cell culture models with both spatial and geometric complexity, we previously introduced Suspended Tissue Engineering with Assemblable Microfluidics (STEAM). STEAM is an accessible, modular platform that utilizes fluidic patterning to create multiregional planar and nonplanar suspended cell-embedded 3D tissues. Herein, we further characterize the STEAM dome platform by developing a theoretical model that explains some experimental considerations necessary for successful two-region patterning in a nonplanar construct. We highlight a brief biological application of the planar and nonplanar STEAM platforms by creating simple but physiologically relevant model systems for the bladder, a sphere-like organ with concentric tissue layers and a central lumen that dynamically expands and contracts during filling and voiding. We demonstrate that the suspended configuration of the planar bladder smooth muscle tissue patch induces inherent tension, which can be increased by further straining the tissues; both result in muscle cell alignment along the axis of stretch as shown by a clear peak at 90° in a radial sum analysis of the two dimensional Fast Fourier Transform of images with fluorescent signal from myosin heavy chain 11 immunostaining. Further, we utilize the nonplanar STEAM platform with a human urothelial cell line (HBLAK) and primary bladder smooth muscle cells (HBdSMC) to create a bladder wall model, resulting in a domed, bilayered tissue. STEAM integrates patterning precision, mechanical functionality, and customizability to actualize an accessible and low cost alternative to generate spatially and geometrically complex suspended tissues.

## INTRODUCTION

Three-dimensional (3D) cell culture, where cells are embedded within a hydrogel network mimicking the extracellular matrix (ECM), advances biological research by enabling more physiologically relevant tissue models in a controlled *in vitro* environment.^1^ Many cell behaviors such as morphogenesis, maturation, proliferation, alignment, and migration are dependent on the mechanical cues provided by the ECM. These mechanical cues include internal cues intrinsic to the hydrogel network such as elasticity and viscoelasticity, as well as external cues applied to the hydrogel network such as compression, stretching, and shear forces.^2,3^ Suspended tissue culture systems, where cell-laden hydrogel constructs are anchored between pillars with variable flexibility, provide cells with mechanical cues from both the ECM and tensile forces generated by the suspended configuration, which can be further modulated through user manipulation of the pillars.^4–7^ These platforms have established their utility for engineering airway muscle,^8^ skeletal muscle,^9,10^ and cardiac muscle tissues^11–13^ by demonstrating the necessity of tension to generate tissues with improved contractile function, alignment, and phenotypic maturity.

Recent work has furthered the physiological relevance of suspended tissue culture systems by incorporating methods for geometric control. Spatial heterogeneity is present throughout biological systems in both diseased (e.g., tumor-stroma interface) and healthy (e.g., bone-ligament interface) tissues. Suspended tissue platforms recapitulate multi-region complexity by incorporating methods for controlled arrangement of cell types and ECM compositions through techniques such as 3D bioprinting^14^ or microfluidic patterning.^15^ Beyond cell and ECM type heterogeneity, cells *in vivo* also experience geometric complexity in the form of nonplanar or curved topography (e.g., enterocytes in colon crypts, epithelial cells in the cornea). Research has shown that cells *in vitro*, both individually and as a population, respond to nanometer-scale surface curvature cues^16^ as well as micrometer-scale cues equal to or greater than cell size.^17–20^ More recently, a few studies have shown that cells cultured on a surface can even sense and react to millimeter-scale (i.e., tissue-scale) curvature. Gouveia and coauthors showed that the presence of a convex agarose substrate in the millimeter range induced corneal stromal cell alignment and increased ECM protein deposition without any additional cues.^21^ Connon & Gouveia demonstrated that concave curvature ranging from 7.5 to 15 mm in diameter increased the migration and alignment of myoblasts parallel to the curvature axis and promoted myoblast differentiation and myotube density.^22^ Several approaches enable 3D nonplanar structures such as casting^23^ and 3D bioprinting.^24–26^ 3D bioprinting can achieve spatial heterogeneity in addition to geometric complexity;^24,27–29^ however, 3D bioprinting often requires complex, custom-engineered systems which can limit accessibility and scalability in some applications.

To provide accessible solutions, we previously developed Suspended Tissue Engineering with Assemblable Microfluidics (STEAM), a platform that generates multi-region planar and nonplanar suspended tissue patches through fluidic patterning.^30^ STEAM employs a disassemblable fluidic channel composed of stackable 3D printed components: a base patterning rail forms the channel floor, a tissue hook device acts as the channel walls, and a top patterning rail serves as the channel ceiling. Cell-laden ECM precursor is loaded into the assembled platform, and the stacked assembly is dismantled after the precursor gels to reveal a tissue suspended between the tissue hook device. The tissue hooks serve as both attachment points and as a mediator for mechanical actuation. The STEAM platform is customizable: the top and bottom patterning rails can be designed to create planar and various nonplanar constructs including wave and dome geometries. In our first introduction of STEAM, we demonstrated the ability of STEAM to pattern a two region dome construct with collagen laden with fluorescently dyed fibroblasts. Here, we show a biologically relevant application of the curvature introduced by the dome construct by creating *in vitro* bladder muscle models in both planar and dome configurations.

The bladder is a mechanically active organ that relaxes during filling and contracts when voiding. Dysregulated contractile behavior of the bladder may lead to overactive bladder symptoms, incontinence or inability to void, or even to elevated bladder pressures leaking to kidney failure.^31^ Most *in vitro* human bladder models are simple organoids or 3D cell culture models designed to study cancer or infection.^32–35^ Organ-on-a-chip approaches incorporate flow for shear strain and urine exposure.^36–38^ However, mechanical function and bladder muscle physiology continue to primarily be studied in small animal models with surgically induced partial bladder outlet obstruction or spinal cord injury.^39–43^ While these *in vivo* models provide the benefit of studying an intact bladder *in situ*, it is difficult to separate direct effects on the bladder from those downstream of the spinal reflexes and autonomic nervous input. In addition, these surgical models are time consuming, costly, and highly variable from animal to animal. Finally, there are known fundamental differences in receptor expression and physiologic response between these animal models and the human urinary tract, limiting the direct translatability of findings from the animal model to patient care.^44^

A recent study from Chae et al bioprinted human bone marrow mesenchymal cells embedded in a decellularized porcine bladder matrix onto a deformable polydimethylsiloxane (PDMS) membrane and demonstrated that cyclical stretch is important for myofibroblast differentiation in 3D culture.^45^ We also know that the bladder urothelium detects mechanical and chemical changes within the bladder and signals both directly and indirectly (via the lamina propria and bladder innervation) to influence the function of the detrusor muscle.^46–49^ With this and other studies demonstrating the importance of spatial and geometric complexity in a variety of muscle models,^50–52^ we decided to apply STEAM to generate an accessible and customizable bladder muscle model with greater biomimicry.

Previously, the STEAM nonplanar multiregion dome device was briefly introduced and tested with dyed mouse fibroblasts as a proof of concept.^30^ In this work, we further characterize the device by examining experimental parameters that affect reproducible placement of multiple regions in a nonplanar construct by developing a theoretical model of capillary pinning in the platform. We then use STEAM to create a bilayered model of the bladder in a nonplanar dome configuration as well as a fixed mechanically strained bladder model with smooth muscle cells in a planar configuration. We demonstrate that spatially and geometrically complex suspended tissues can be achieved reproducibly through STEAM, thus showcasing STEAM’s strength to model curved organs in the body such as the bladder.

## RESULTS AND DISCUSSION

### STEAM is an accessible, modular platform that can generate multiregional, nonplanar tissue constructs

STEAM is a modular platform that utilizes removable, stackable 3D printed parts to form a microfluidic channel that enables patterning of complex, three-dimensional suspended tissues using a standard pipette. This channel is formed by a base patterning rail, a tissue hook device that also forms the channel walls, and a top patterning rail; which together assemble into the “floor,” “walls,” and “ceiling” of a temporary fluidic channel, as described in detail previously (Figure 1ai and 1aii).^30^ Because tissue shape is dictated by the shape of the patterning rails, this same assembly principle extends to a broad range of construct geometries, from flat planar patches to complex nonplanar architectures, simply by changing the rail design.^30^

**Figure 1:**
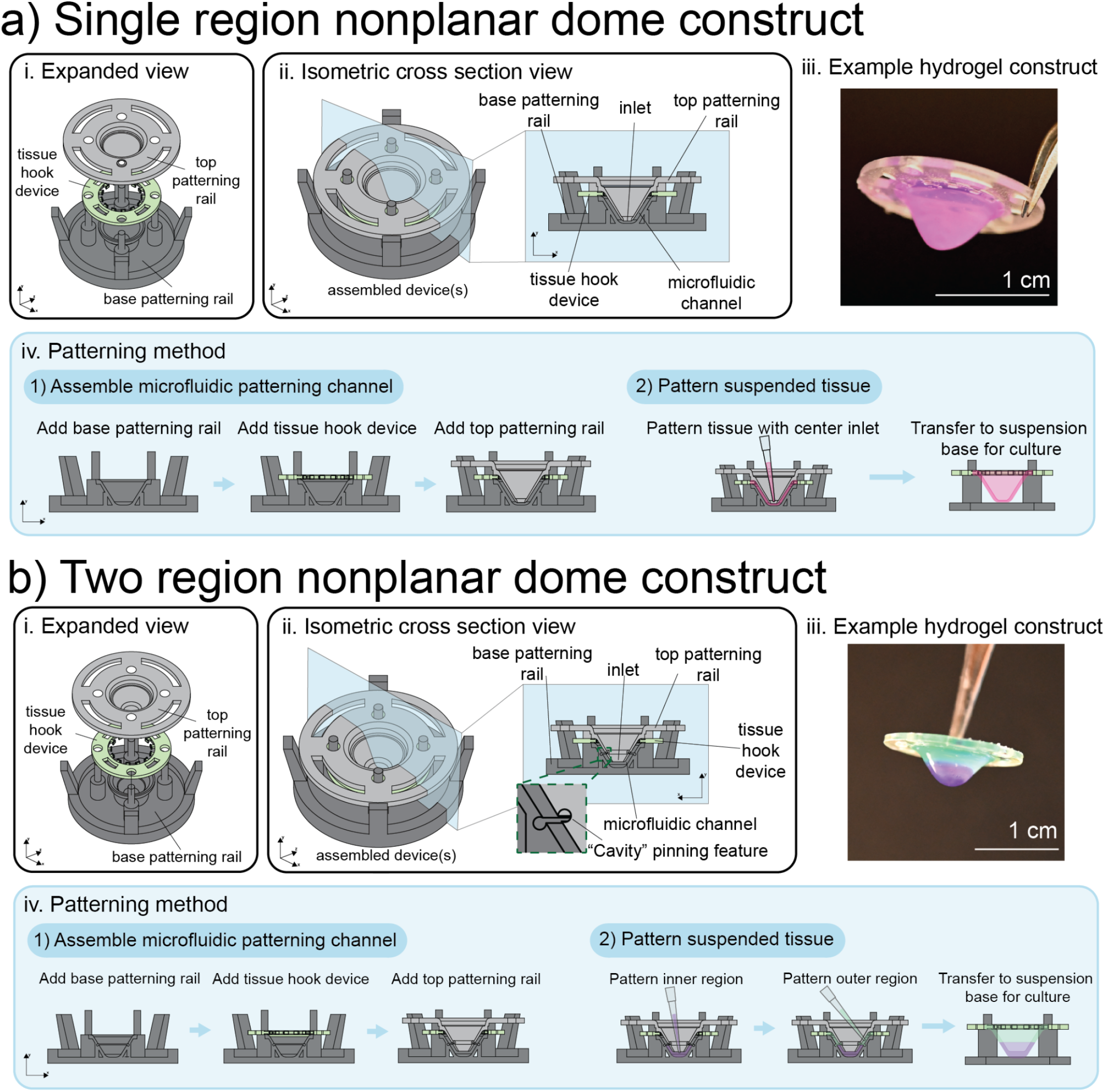
Suspended Tissue Engineering with Assemblable Microfluidics (STEAM) as a customizable platform for tissue generation. **a)** The single region nonplanar dome STEAM platform consists of a base patterning rail, tissue hook device, and top patterning device (i). When assembled, the stack forms a microfluidic channel (ii) capable of generating a nonplanar dome construct as demonstrated with dyed collagen (iii). The platform construction occurs as follows: first, a base patterning rail is placed in a 6-well plate, a tissue hook device is layered on top, and finally, the top patterning rail is stacked above for the full assembly (iv, step 1). A simple pipette filled with hydrogel precursor can then be used to pattern a tissue from the center inlet of top patterning rail (iv, step 2). After gelation time, the top patterning rail can be removed, and the construct can be moved to a suspension base for long term culture (iv, step 2). **b)** The two region nonplanar dome STEAM platform consists of a base patterning rail, tissue hook device, and top patterning device (i). When assembled, the stack forms a microfluidic channel with a “Cavity” pinning feature revolved radially in the channel (iii), which allows for two-region, contiguous dome tissue generation as seen in an example image of patterned collagen (iii). The patterning method begins by assembling the devices as described above (iv, step 1). After the devices are assembled, the inner region is patterned by dispensing hydrogel precursor in the center inlet first, followed by dispensing hydrogel precursor in flanking inlet(s) for the outer region (iv, step 2). After gelation time, the top patterning rail can be removed, and the construct can be moved to a suspension base for long term culture (iv, step 2). All scale bars are 1 cm.

Here, we focus on one such nonplanar geometry, a dome-shaped construct, motivated by its relevance to modeling curved tissue surfaces in the body such as the bladder wall. Unlike planar STEAM or string-like constructs, which are elongated between two discrete suspension points,^11,15,53–57^ the dome construct is radially symmetric, producing a curved tissue (Figure 1aiii). The patterning process begins by assembling the STEAM dome platform with the following steps: first, a base patterning rail is inserted into a 6-well plate; then, a tissue hook device, that serves as anchor points for the tissue, is placed on top of the base patterning rail; finally, a top patterning rail is stacked on top of the assembly (Figure 1aiv, step 1). The suspended nonplanar dome construct is then made by pipetting hydrogel precursor into the center inlet of the top patterning rail where the precursor flows radially up the channel to surround the tissue hook device anchors (Figure 1aiv, step 2). The tissue hook device forms a continuous rim around the circumference of the dome; this radial symmetry creates suspension along the construct’s full perimeter.

Like in previous planar demonstrations of two-region patterning,^15,30^ we utilize a concave “cavity” pinning feature that locally halts the advancing fluid front of the first pipetted region, creating a stable boundary against which a second region can then be pipetted to form a contiguous construct without the two regions mixing (Figure 1bi and ii). In the dome configuration, the pinning feature runs circumferentially within the channel, partitioning the dome into two concentric regions, a central inner region and a surrounding outer region, that meet at a circular rather than linear boundary (Figure 1bii). Representative dome constructs patterned with colored collagen hydrogel solutions show a well-defined, concentric boundary between regions following transfer to a suspension holder for culture (Figure 1biii). Two-region patterning of the dome construct follows the same underlying workflow established for planar STEAM patches: after assembling the channel (Figure 1biv, step 1), the inner region is pipetted first and confined by the cavity pinning feature; then, the outer region is pipetted to fill the remaining channel volume up to the pinned interface, forming a contiguous two-region construct; finally, the patterned construct is transferred to a suspension base for longer term culture after the hydrogel precursor solidifies (Figure 1biv, step 2). Notably, nonplanar STEAM generated tissues can also be utilized for bilayered constructs by seeding cells suspended in liquid media in the center of the dome. These bilayered constructs can better mimic certain coculture environments in the human body such as organs that comprise multiple heterogeneous tissue layers.

### Theoretical modeling of the capillary pinning mechanism in nonplanar STEAM

The STEAM dome platform utilizes a cavity pinning feature that spans the circumference of the platform to achieve two region patterning (Fig 2a). Previous work has theoretically modeled and experimentally validated the mechanism for successful capillary pinning by the cavity pinning feature in our previously published Suspended Tissue Open Microfluidic Patterning (STOMP) platform, which features a string-like, suspended channel that lacks both a channel floor and a channel ceiling.^15^ Since the channel in the STEAM dome platform is non-planar and semi-open (i.e., the channel is enclosed on all sides except at the inlet), different factors affect the success of capillary pinning by the pinning feature and a new theoretical model is needed. Here, we advance the theoretical knowledge of parameters affecting capillary pinning in nonplanar, semi-open systems by examining the effect of hydrostatic pressure from the fluid in the pipette and fluid escape from the inlet on the ability of the pinning feature to arrest flow.

**Figure 2:**
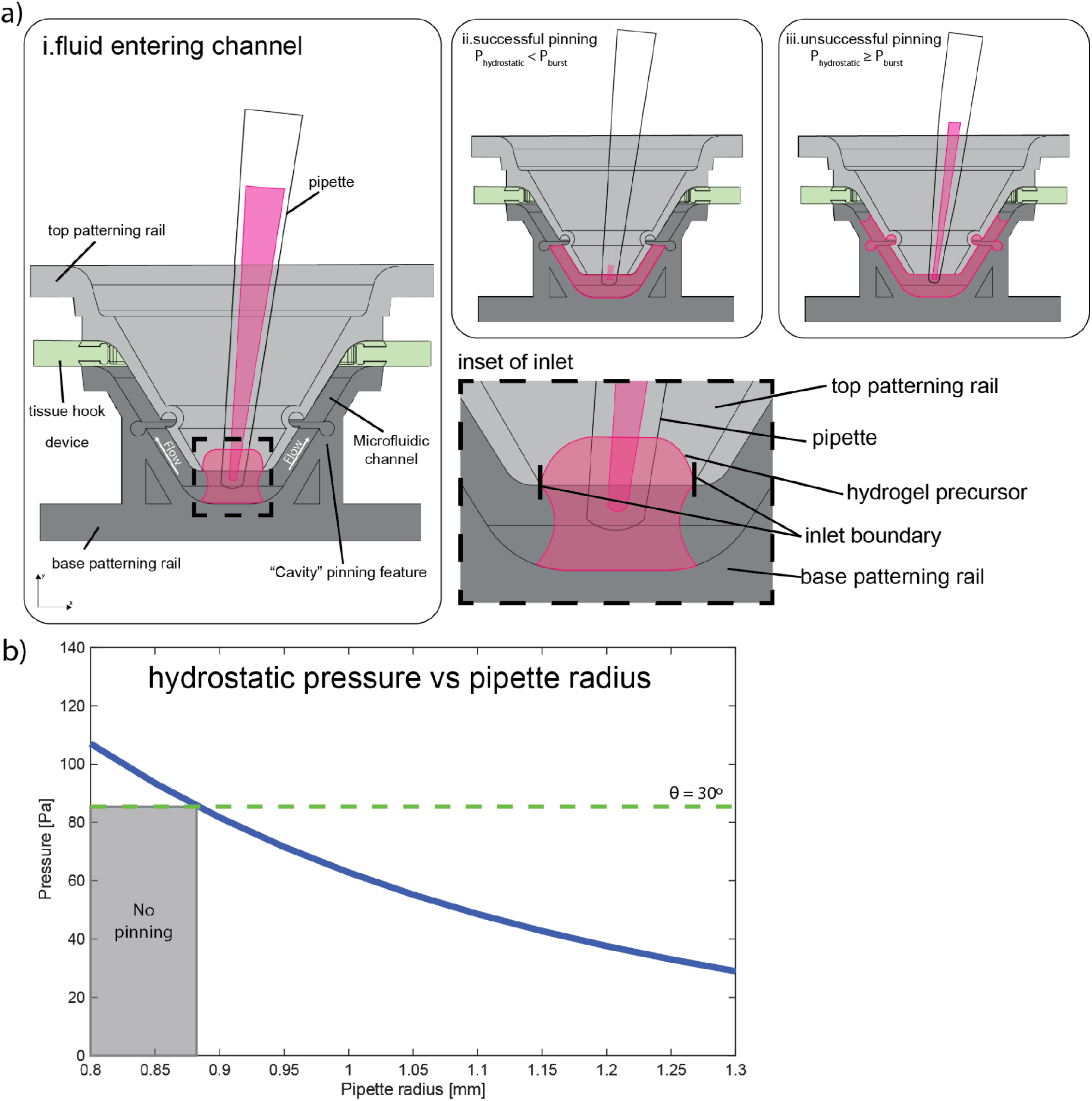
Characterization of design considerations to generate multiregional nonplanar STEAM constructs. **a)** Schematic depicting the mechanism for pinning success and failure in the STEAM dome platform. Hydrogel precursor is pipetted into the channel, the precursor fluid front advances along the channel (i) with some of the precursor escaping at the channel inlet (inset). In a scenario with successful patterning, the height of the residual fluid in pipette after dispensing the precursor is low, the hydrostatic pressure (*P*_ℎ*ydrostatic*)_ is less than burst pressure (*P_burst_*), and the fluid front remains pinned by the cavity pinning feature (ii). In a scenario with unsuccessful patterning, the height of the remaining fluid is high, the hydrostatic pressure (*P*_ℎ*ydrostatic*_) is equal to or greater than burst pressure (*P_burst_*), and the fluid front pushes past the pinning feature. **b)** Graph of hydrostatic pressure plotted against pipette radius based on the derived theoretical model. The green dashed line corresponds to the burst (depinning) pressure for a liquid with a contact angle (θ) of 30° pinned at the cavity pinning feature. For fixed total volume of fluid, hydrostatic pressure and pipette radius have an inverse relationship. Depinning is predicted to occur when the pressure associated with the pipette radius is greater than the burst pressure for a given fluid and capillary pinning feature, which is shaded in gray on the graph.

Passive fluid flow in microfluidic channels is generally governed by Laplace pressure, the pressure difference between inside of the fluid and outside of the curved fluid front (i.e., the advancing meniscus in the channel), and hydrostatic pressure, the pressure exerted on a fluid at rest due to gravity. After the hydrogel precursor is displaced from the pipette during the patterning process, a small portion of hydrogel precursor remains in the pipette tip which contributes hydrostatic pressure to the system. In the STOMP platform, the channel is fully open, the inlet is markedly larger than the pipette tip, and the pipette is generally withdrawn before the fluid front reaches the pinning feature.^15^ Thus, the hydrostatic pressure presented by residual fluid in the pipette tip is minimal. In contrast, the STEAM dome platform has a semi-open channel and the pipette tip remains within the channel throughout the patterning process (Figure 2ai). Therefore, the hydrostatic pressure from the remaining fluid in the pipette tip is an important consideration for the theoretical model of the capillary pinning mechanism in the STEAM dome platform. When the fluid front reaches the cavity pinning feature, hydrostatic pressure is given by the equation

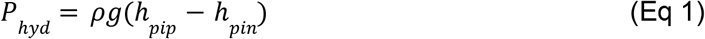

where *ρ* is the density of the fluid, *g* is acceleration due to gravity, ℎ*_pip_* is the height of the fluid column in the pipette, and ℎ*_pin_* is the height of the pinned interface. Since the remaining fluid in the pipette tip is the source of the hydrostatic pressure in the STEAM dome platform, the varying height of the fluid in the pipette tip is the main determinant of the amount of hydrostatic pressure present in the system. When pinning is established, the Laplace pressure of the pinned interface balances the hydrostatic pressure. Additionally, we experimentally observe some of the hydrogel precursor escaping upwards from the channel inlet (Figure 2a inset) due to the small open gap between the pipette and the inlet. The escaping precursor forms a fluid ring around the pipette at the channel entrance where the interface of the fluid ring is defined by contact with the pipette exterior and anchoring to the top surface of the inlet edge. The escaping precursor reduces the volume in the pipette—and thus height of the residual fluid in the pipette—which decreases the hydrostatic pressure introduced to the system by the fluid in the pipette.

In a successful pinning scenario, after dispensing the hydrogel precursor into the channel via pipette, the fluid front advances in the channel until it reaches the cavity pinning feature. The fluid front then halts as the pinning feature presents an abrupt enlargement of the channel cross-section that is energetically unfavorable to overcome (Figure 2aii).^58,59^ The amount of pressure required to drive the fluid front past the pinning feature is known as the burst pressure, which is dependent on the contact angle of the fluid and the geometry of the pinning feature.^60^ For successful pinning, the hydrostatic pressure must not exceed the burst pressure. Once hydrostatic pressure is equal to or greater than the burst pressure, depinning occurs (Fig 2aiii). A detailed derivation for the relationship between the hydrostatic pressure and the burst pressure in the STEAM dome platform can be found in the Supporting Information.

By creating a theoretical model that relates hydrostatic pressure and burst pressure, as detailed in the Supporting Information, we identified the radius of the pipette as a key parameter that determines successful capillary pinning (Figure 2b). For a given total volume, when the pipette radius decreases, the height of the fluid in the pipette increases, proportionally increasing the hydrostatic pressure. Once the hydrostatic pressure associated with a pipette radius exceeds the burst pressure, which is dependent on the contact angle of the pinning fluid and represented by the green dashed line, depinning occurs. The area highlighted in red shows where pinning is predicted to fail. In this graph, the total volume is fixed and pipette radius is varied. Using the relationship between fluid volume and fluid height, a similar relationship could be plotted for varying total fluid volume for a fixed pipette radius. Thus, pipette radius and total fluid volume are both parameters that govern successful capillary pinning. Furthermore, the relation between the pipette radius and the inlet radius of the top patterning rail determines the size of the gap from which fluid escapes the semi-open channel and forms a fluid ring around the pipette. Since the fluid ring may aid in pinning success by decreasing hydrostatic pressure presented by the fluid remaining in the pipette, future work will model the effects of different inlet radii for a fixed pipette radius on the success of capillary pinning with additional studies on dome systems of different dimensions such as height. Altogether, our theoretical model provides a new understanding of the mechanism behind capillary pinning in nonplanar, semi-open systems and highlights potential parameters that should be considered for successful pinning.

### STEAM allows exertion of variable strain on bladder smooth muscle tissues

Since many tissues, including the bladder detrusor and other muscles, need mechanical cues for proper development and function,^45,61–63^ STEAM creates suspended tissues with inherent tension and leverages the tissue hooks to act as anchor points for adherence during suspended culture and downstream mechanical strain application. In planar, STEAM generated constructs for mechanical strain application, the tissue hook devices are two independent components where strain is applied along a single axis by moving the independent tissue hooks separately to a set amount of static strain (Figure 3a-d). Each set of hooks contain a peg on the bottom which interfaces with a corresponding hole in an incremental stretching device, allowing for a planar patterned tissue to be to be held in suspension at a particular amount of strain. Therefore, a transfer device was needed in order to stabilize the tissue during transfer from the base patterning rail to the incremental stretching device. In the present work, we introduce an updated transfer device building upon the device used in Whitten & Haack.^30^ Instead of a press-fit feature that depended on 3D printer accuracy and reproducibility on a small scale, a snap-fit was integrated to join the tissue hook device(s) with the transfer device (Figure 3a, device shown in blue and Figure 3b) for a facile transfer of the two independent tissue hook devices in tandem from the base patterning rail. Following transfer to the incremental stretching device, the bottom pegs of the tissue hook device(s) secure the hook device in a predefined position. Controlled strain is then applied by sequentially repositioning the tissue hooks one at a time using tweezers into holes on the incremental stretching device spaced at predetermined distances (Figure 3c-d). This allows for the tissue to achieve a stretch up to a 5 mm increase (50% strain for the 10 mm tissue patch); however, for the purpose of this manuscript, only a stretch up to a 2.5 mm increase (25% strain for the 10 mm tissue patch) was used for the bladder smooth muscle model per previous literature.^64–66^

**Figure 3:**
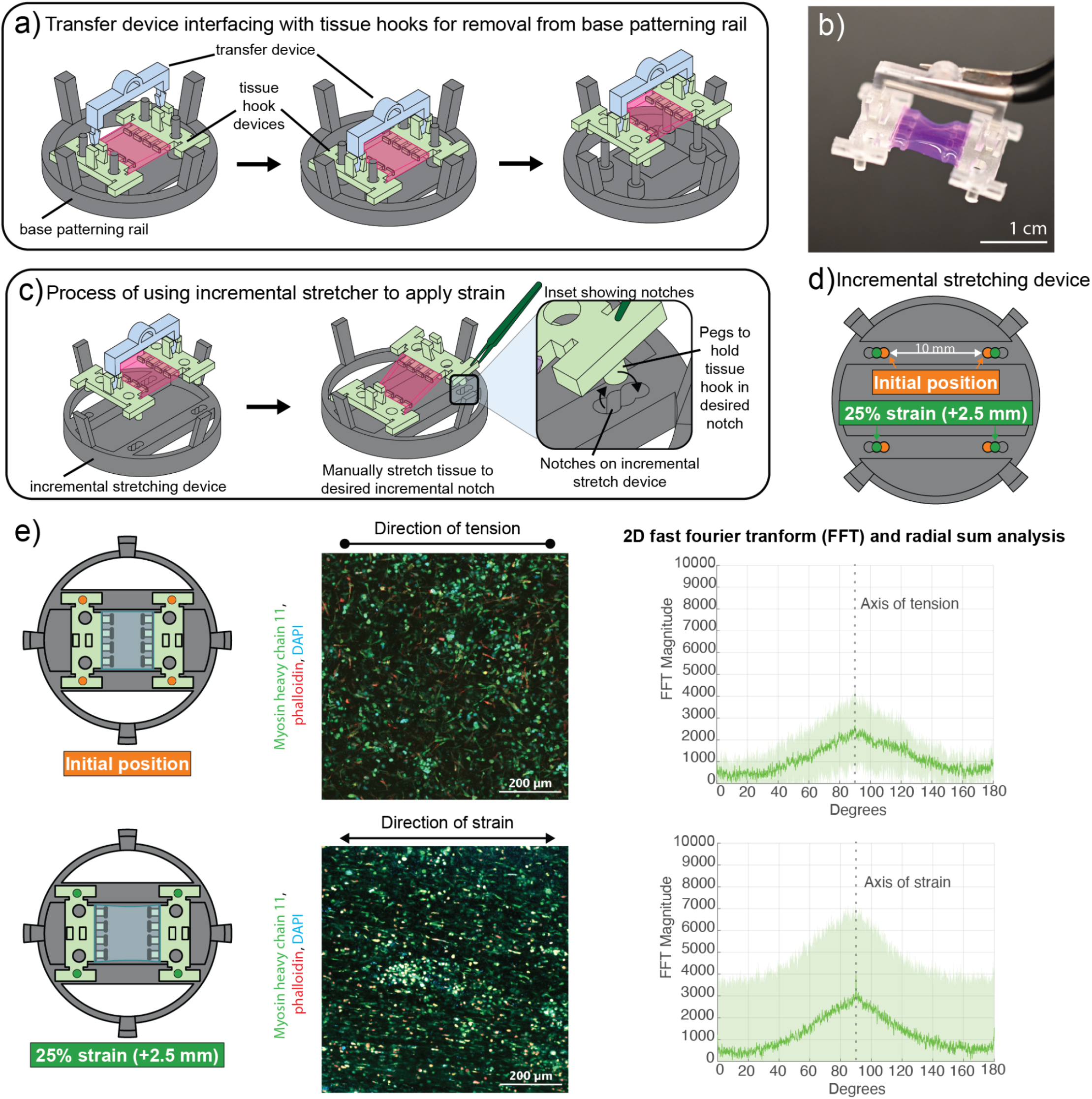
STEAM allows exertion of variable strain on bladder smooth muscle tissues. **a)** Schematic workflow for lifting a tissue off the patterning setup using the transfer device. The transfer device uses a snap-fit feature to temporarily attach, and stabilize, the tissue hooks for transfer from the base patterning rail. Recreated from Whitten & Haack.^30^ **b)** An example photo of the transfer device being used to lift a dyed collagen planar patch with tweezers. Scale bar is 1 cm. **c)** Schematic workflow of placing the tissue on the incremental stretching device **(d)** by aligning the pegs on the bottom of the tissue hook devices with the “initial position” notches on the incremental stretching device. Next, the tissues can be manually stretched by lifting the tissue hook device and moving it so the pegs are placed in a different notch, inducing a set strain. Recreated from Whitten & Haack.^30^ **e)** Schematic showing the initial position (top) and 25% strain (bottom) tissues as a top-down view on the incremental stretching device followed by a representative confocal image of tissues after 10 days of culture and post-fabrication applied strain. The confocal images show a maximum projection of HBdSMCs stained for the nucleus (DAPI, blue), cytoskeleton (phalloidin, red), and a bladder smooth muscle cell marker (myosin heavy chain 11, green). Scale bar is 200 µm. 2D fast fourier transform (2D FFT) and radial sum analysis of the two experimental conditions, with shading representing standard deviation of three regions of interest in a single tissue. 90° is defined as the direction of stretch as peak frequency is calculated perpendicular to the direction of stretch in a more aligned tissue.

As described previously, the bladder exemplifies why tissue-scale mechanical modeling matters. During each fill-void cycle, the bladder’s multiple tissue layers—each exhibiting distinct mechanical properties—experience extreme mechanical deformation.^67^ Therefore, the STEAM platform is well suited to demonstrate the functional impact of applied mechanical manipulation to tissue response by investigating induced alignment of primary human bladder smooth muscle cells (HBdSMCs) embedded in collagen when placed under a set strain. We first verified that the HBdSMCs expressed a well established bladder smooth muscle cell marker, myosin heavy chain 11 (MYH11), which is shown in the immunofluorescence images of both the control and the 25% strain tissues (Figure 3e).

We hypothesized that applying added strain (25%) to a suspended tissue with embedded HBdSMCs would induce increased alignment of the cells along the axis of strain in comparison to HBdSMCs solely in suspension. To visualize cellular structures, tissues after 10 days of post-fabrication stain were immunostained with MYH11 (Figure 3e). Utilizing a radial sum analysis of the two-dimensional fast fourier transform (2D FFT) of confocal fluorescent images as previously described^30,68^, we saw an increase in HBdSMC alignment from the control (0% added strain) in comparison to 25% strain (Figure 3e). With regards to the radial sum analysis, the degree of cellular alignment in an image is reported by the height and shape of the peak generated by the radial sum of the 2D FFT plot across 180° at a specified radius. Therefore, if MYH11 were randomly oriented in the tissue, there would be no peak at any angle on the 2D FFT plot. Broader peaks indicate moderate alignment to an angle with some unaligned fibers, and high peaks indicate a uniform degree of alignment. We note the peak shape in the control tissue reflects some degree of alignment based on the singular peak oriented at 90° which is the direction of tension in our system; this is due to the tissue being held in a suspended configuration which adds some degree of inherent tension. Based on the 2D FFT plot of our control (0% added strain) and the 25% strain tissues, the 25% strain tissue trended towards more alignment in comparison to the control (Figure 3e), with a 49% increase in peak height. However, we note the variability between replicate images used to generate the 2D FFT plot is high in our initial experiment, and current work is under way to compare different degrees of strain for varied amounts of time.

### Utilization of STEAM to create a nonplanar, bilayered bladder model

The bladder wall comprises three main layers: the urothelium, the lamina propria, and the muscularis propria, each with specific functions.^69,70^ Noting the bladder is a sphere-like structure with concentric tissue layers, we used STEAM to model a hemisphere (dome) to partially recapitulate the curvature seen with the bladder. We first patterned collagen precursor laden with HBdSMCs in the STEAM dome platform (Figure 4a). Once the collagen gelled, the top patterning rail was removed and urothelial cells (HBLAK) suspended in liquid media were pipetted inside the dome structure and left to adhere. After the HBLAK cells adhered, the tissue construct was then removed from the base patterning rail and cultured in suspension to overall emulate the concentric tissue configuration present in the bladder wall. Tissues were left in culture for 4 days before fixation and subsequent tissue sectioning. Tissue architecture was validated with hematoxylin and eosin (H&E) staining, a widely used set of dyes that provide a contrasting view of cell structure and tissue architecture where the cell nucleus is stained purple and the surrounding area—which can be the cytoplasm, collagen, or other ECM—is stained pink. The H&E stain of the dome tissue section demonstrates the successful creation of a bilayer where cells are both embedded in the collagen matrix and adhered on top of the collagen (Figure 4b). Ongoing work includes longer tissue culture durations, more detailed characterization of cellular differentiation and organization, and using the radial sum analysis of a 2D FFT plot in a nonplanar, suspended dome construct to assess the effect of curvature on cellular alignment.

**Figure 4:**
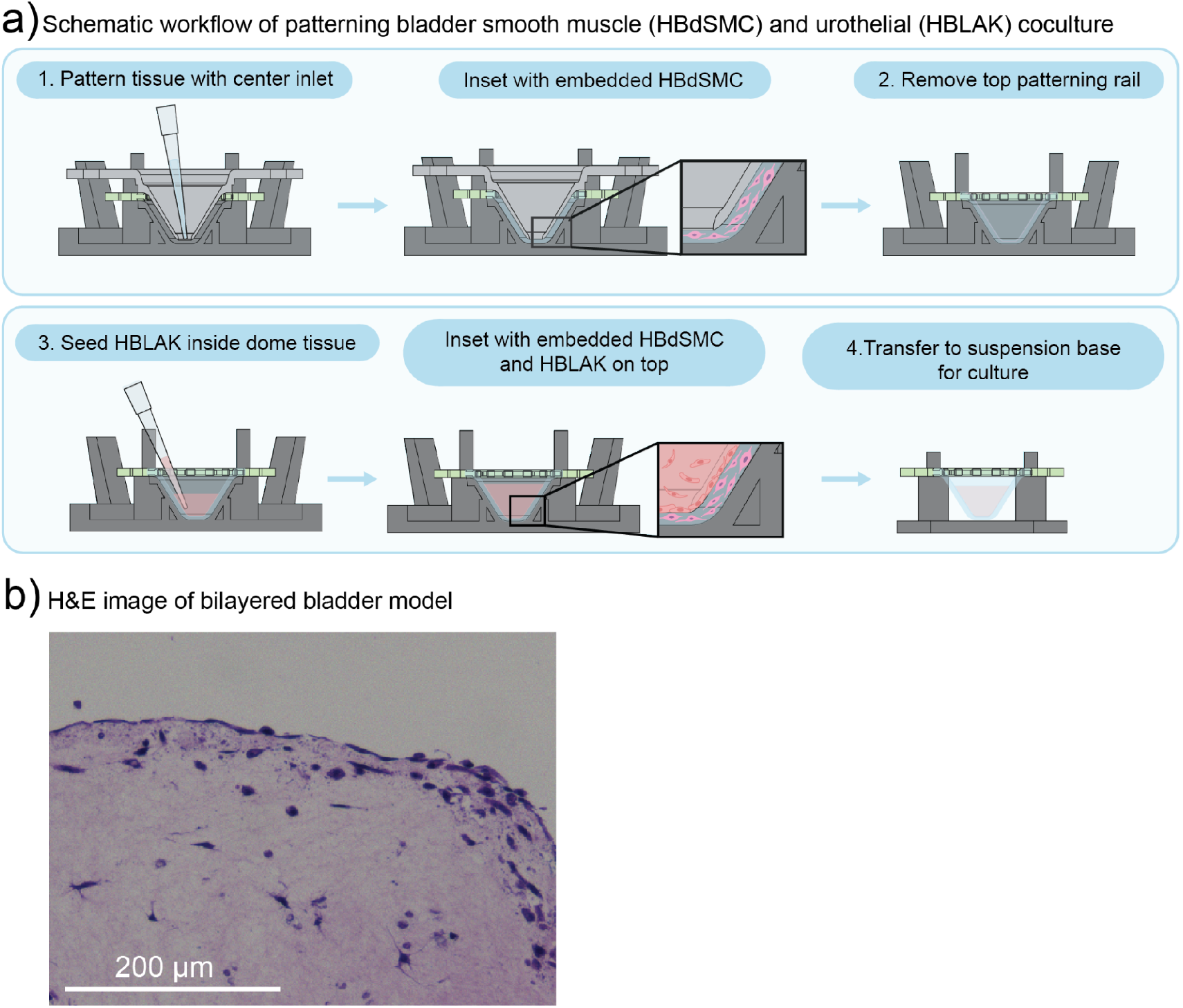
Creating a nonplanar, bilayer bladder model with STEAM. **a)** Schematic workflow for coculture setup. First, HBdSMC-laden hydrogel precursor is patterned with the dome platform (1) and incubated at 37 °C to gel. Inset shows the embedded HBdSMC in the platform. Next, the top patterning rail is removed from the patterning scheme (2) and HBLAK cells suspended in liquid media are seeded on the concave surface of the HBdSMC-embedded dome tissue (3). Inset shows after HBLAK cells are left to adhere on top of the tissue with embedded HBdSMC. Finally, the tissue construct can be transferred to a suspension base within a 12-well plate for long term culture (4). **b)** H&E stained image of tissue section after 4 days in culture. The collagen matrix is shown in pink with cells stained purple. Cells are both embedded in the collagen matrix and adhered to the exterior of the collagen structure. Scale bar is 200 µm.

## CONCLUSIONS

In this paper, we further developed and validated STEAM as a customizable platform to model nonplanar and mechanically complex tissues. STEAM employs modular, removable fluidic channel elements that define customizable spaces for tissue fabrication to expand the geometric and mechanical possibilities available for suspended tissue engineering. By leveraging microfluidic principles, a theoretical model based on hydrostatic pressure was developed to explain various experimental considerations important for reproducible multiregional patterning in a nonplanar tissue construct mediated by capillary pinning. Simple models of the bladder were created to demonstrate STEAM’s ability to generate curved architectures that are ubiquitous in biology, as well as its potential to create models with tunable mechanical properties to enable direct study of the physiologically relevant mechanobiology within tissue constructs. Future extensions may incorporate multiple pinning features to increase tissue interface complexity as well as an expansion to multiregionality on the surface of a tissue construct.

## METHODS

### Device design and fabrication

All STEAM device patterning components were designed in SolidWorks and 3D printed out of clear V4.1 resin using Form 3/3B stereolithography 3D printers (FormLabs Inc.). Devices underwent a standard cleaning sequence to remove excess uncured resin. This sequence begins with agitation in isopropyl alcohol (IPA) for 20 min followed by a second agitation for 10 min in two separate FormWash units (Formlabs Inc.). Devices were then transferred to beakers filled with fresh IPA for 30 min of sonication, after which devices were dried with pressurized air or set out on the bench until dry. Post-drying, devices were post-cured with 405 nm light at 60 °C for 15 min in a FormCure (Formlabs Inc.). Prior to biological experiments with cells that involve stretching the tissues, the tissue hook devices were treated by oxygen plasma using a Diener Zepto PC EX Type PB plasma treater (Diener Electronic, Germany). The plasma treatment improves tissue adherence to the tissue hooks during stretching and subsequent culture. All device components were exposed to UV light for a minimum of 15 min in the biosafety cabinet (BSC). After UV exposure, the top and base patterning rails were incubated in a solution of 1% bovine serum albumin (BSA) for 1 h at room temperature to aid in tissue removal and achieve appropriate SCF conditions. After incubation, the 1% BSA solution was aspirated, and the patterning rails were allowed to fully dry prior to assembling for patterning.

### Patterning one- and two-region hydrogel STEAM constructs with collagen

The top and base patterning rails for both one- and two-region STEAM devices were incubated in a solution of 1% BSA for 1 h at room temperature. After incubation, the 1% BSA solution was aspirated and the patterning rails were allowed to fully dry prior to assembling. One- and two-region STEAM patterning devices were assembled in the following order: first, the base patterning rail was placed in a 6-well plate; then, the tissue hook device was placed onto the base posts; finally, the top patterning rail was placed on the stacked assembly. The collagen solution was prepared by diluting a stock of rat tail collagen type I in 0.02 N acetic acid (Corning) with deionized water, 10X phosphate buffered solution (PBS), 1M HEPES buffer solution, 7.5% sodium bicarbonate solution, and 1 N sodium hydroxide (NaOH) to achieve a final collagen density of 5 mg mL^-1^ and kept on ice to prevent gelation. The deionized water was colored with food dye (Spice Supreme) prior to collagen solution preparation to aid in visualization (Images shown in Figure 1). For a one-region, nonplanar 6 mm height cone device 210 µL of collagen was pipetted into the center inlet and incubated for 30 min at 37 °C for gelation. For a two-region, nonplanar 6 mm height cone device, 55 µL of collagen is pipetted into the center inlet for the inner region, followed by 147 µL of collagen in the outer inlet(s) for the outer region. The two-region, nonplanar cone device was allowed to gel for 45 min at 37 °C before 1X PBS was added to the 6-well plate to facilitate top patterning device removal. After device disassembly, the tissue hook device with the collagen construct was transferred to a separate tissue holder in a 12-well plate and submerged in 1X PBS. Images of collagen constructs were taken using a Nikon D5300 DSLR high resolution camera.

### Cell culture maintenance - HBLAK

Spontaneously immortalized, nontransformed human bladder epithelial cells, HBLAK, were obtained from CELLnTEC. The cells were maintained in tissue culture flasks containing PneumaCult-NGEx proliferation media with 50X supplement (StemCell Technologies Inc.) at 37 °C, 5% CO_2_. Culture medium was changed every 48h until cells reached 80-90% confluency, whereupon the culture was rinsed with 1X PBS, followed by addition of Accutase (Sigma-Aldrich). After incubation for 6-8 min at 37 °C, the Accutase was inactivated by diluting with cell culture medium at a 2:1 ratio. The fluid volume was centrifuged at 200 RCF for 5 min to pellet the detached cells. The cells were then resuspended in cell culture medium for further passaging or used for patterning experiments described below. HBLAK cells between passage numbers 3 and 7 were used for patterning experiments.

### Cell culture maintenance - HBdSMCs

Primary human bladder smooth muscle cells (HBdSMC) were obtained from ATCC and maintained following ATCC protocols. Briefly, the cells were maintained in tissue culture flasks containing vascular cell basal medium with a vascular smooth muscle cell growth kit (ATCC) at 37 °C, 5% CO_2_. Culture medium was changed every 48h until cells reached 80% confluency, whereupon the culture was rinsed with 1X PBS, followed by addition of TrypLE Express (Gibco). After incubation for 3-5 min at 37 °C, the TrypLE was inactivated by diluting with cell culture medium at a 2:1 ratio. The fluid volume was centrifuged at 150 RCF for 5 min to pellet the detached cells. The cells were then resuspended in cell culture medium for further passaging or used for patterning experiments described below. HBdSMCs between passage numbers 4 and 9 were used for patterning experiments.

### Patterning and stretching HBdSMCs in collagen with the static stretch device

After dissociating the HBdSMCs, they were pelleted and resuspended at a final concentration of 4×10^6^ cells mL^-1^ in a liquid collagen solution described above. Static stretching STEAM devices were assembled in a 6-well plate and 230 µL of cell-laden ECM-precursor was pipetted in the inlet and allowed to gel for 45 min at 37 °C. After gelling, the top patterning rail was removed and the transfer device was slid into place on the tissue hook devices to aid in controlled removal. The transfer device was then used to move the tissue hooks in tandem to the incremental stretching device in another 6-well plate. After transfer, 6 mL of media was added to each well. After 24 h in culture, tissues were randomly selected for controls (0% strain) and 25% strain. The tissues selected for increased strain had the tissue hooks moved from the control configuration to the 25% strain configuration with tweezers. Tissues were cultured for an additional 10 days with media refreshed every 48 h.

### Coculture of HBdSMCs and HBLAKs in nonplanar cone device

After dissociating the HBdSMCs, the cells were pelleted and resuspended at a final concentration of 4×10^6^ cells mL^-1^ in a liquid collagen solution as described above. 6 mm height nonplanar dome STEAM devices were assembled in a 6-well plate, and 230 µL of cell-laden ECM-precursor was pipetted in the inlet and allowed to gel for 45 min at 37 °C. After gelling, 1X PBS was added to the well plate to aid in removal of the top patterning rail. The tissue hook device was then transferred to a separate tissue holder in a separate 12-well plate for longer term culture. The HBLAKs were then dissociated, pelleted, and resuspended at a final concentration of 3×10^5^ cells mL^-1^ in HBLAK associated media. Approximately 2 mL of HBdSMC associated media was added to each well of the 12-well plate containing a suspended dome with embedded HBdSMCs in collagen to provide appropriate nutrients to the HBdSMCs while 100 µL of HBLAK cell suspension was added to the top of the tissue construct, filling the dome structure. HBLAK cells were left to adhere for approximately 2 h at 37 °C before the tissue was submerged in a 3 mL of 1:1 ratio of HBdSMC and HBLAK media. Tissues were cultured for an additional 4 days with media refreshed every 48 h.

### Immunofluorescence staining and imaging

Tissue constructs were rinsed twice with 1X PBS and fixed at 4% PFA at 4 °C for 1 h. Tissues were then rinsed three times with 1X PBS and stored at 4 °C until immunofluorescence (IF) staining. For IF staining preparation, the tissues were permeabilized with 0.5% Triton-X (v/v) for 1 h followed by blocking with 10% FBS for 3 h at room temperature with gentle agitation from a plate shaker set at 200 RPM. Tissues were then incubated with a primary antibody overnight at 4 °C, washed with a 0.2% Triton-X (v/v) solution, and incubated with a secondary fluorescent antibody, Alexa Fluor 647 Phalloidin (Molecular Probe) (1:100), and 4′,6-diamidino-2-phenylindole (DAPI) (Molecular Probe) (1:500) overnight again at 4 °C. Tissues were washed with 1X PBS before imaging. The primary antibody used was anti-smooth muscle myosin heavy chain 11 (abcam, ab133567) (1:50) and the secondary antibody used was Alexa Fluor 488 goat anti-rabbit (Jackson ImmunoResearch) (1:100). Images were acquired using a 20X magnification on a Zeiss LSM 900 confocal microscope equipped with ZEN software 3.0. The confocal images obtained were split into individual channels per fluorophore used for image analysis with ImageJ v1.54t or later and MatLab 2024b. Confocal images displayed in Figure 3e were adjusted in ImageJ v1.54t on a representative region, increasing brightness and contrast, and applying a Gaussian filter (σ=2) to reduce noise uniformly within and across images.

### HBdSMC static stretch patch alignment quantification image analysis

NIH imageJ software was used to analyze the alignment of myosin heavy chain 11 within control (0% added strain) planar HBdSMCs tissues and planar HBdSMC tissues placed under 25% strain in the static stretch device, as previously described.^30,68^ Briefly, confocal images were loaded into MatLab and downsized to 7434 by 7434 pixels using bicubic interpolation to dimensions compatible with the NIH imageJ radial sum macro developed in-house. The 2D fast fourier transform (2D FFT) of the green (MYH11) channels of the confocal image were taken for analysis. The 2D FFT converts the spatial data of the image into a mathematically defined frequency domain. The resulting plot has equivalent dimensions to the original image in which the center encodes the magnitudes of low-frequency image features and the outer region encodes the magnitudes of high-frequency features, while the angle represents the direction of the frequency in the spatial image. On each 2D FFT, an oval region of interest of 2500 by 2500 pixels, used in previous work to enclose the region of frequencies representative of myosin heavy chain patterns, was plotted in imageJ using an oval profile plugin. A radial sum was taken at 3000 points along the oval region of interest, and these summed magnitudes were plotted against their corresponding angle. As is intrinsic to its computation, the 2D FFT plot is transformed 90° from the orientation of the spatial image, so peaks at 90° represent frequencies aligned with 0° in the image, which is the axis of strain. 2D FFT and radial sum analysis was run with ImageJ v1.54t, and image resizing was done in MatLab 2024b.

To perform the above method, confocal max projection images of three representative regions of interest per single tissue were used. Within images and across all images, brightness was uniformly adjusted in ZEN3.0 Microscopy Software before FFT analysis. Radial sums of the three regions of interest were averaged, normalized, and plotted in MatLab 2024b. Radial sum plots were normalized to a baseline value of 0 and plotted in arbitrary units, allowing different plots to be directly compared, resulting in the magnitudes plotting along the y-axis of Figure 3e. All values are reported as mean ± standard deviation across the three regions of interest. Percent change in peak height was calculated using the maximum magnitude of the control (0% added strain) versus the 25% strained 2D FFT magnitude plot.

### H&E staining

Tissue constructs were rinsed twice with 1X PBS and fixed at 4% PFA at 4 °C for 1 h. Tissues were then rinsed three times with 1X PBS and stored at 4 °C. Tissues were paraffin embedded and sectioned at a thickness of 4 µm by the UW MANTIS Lab (core facility). Hematoxylin and eosin (H&E) were then used to stain the sections in standard fashion by the UW MANTIS Lab. Images were acquired with a BioTek Cytation 5 plate reader (Agilent) with a 20X objective on the colored bright field setting.

## Acknowledgments

This publication was supported by the National Institutes of Health (NIH) through the National Institute of General Medical Sciences award number R35GM128648 (ABT). This publication was also partially supported by an American Urologic Association Research Scholar Award (MRC). The content is solely the responsibility of the authors and does not necessarily represent the official views of the National Institutes of Health or other funding bodies.

## Author Contributions

JMW, EEB, EB, ABT, AJH, and MRC conceived the project. EB, ABT, AJH, and MRC supervised the project. JMW and EEB designed and fabricated the device with process inputs from EAS, ABT, and AJH. JB developed the theoretical model for flow and pinning in the two region system with inputs from JMW and AL. JMW, EEB, EAS, and AL designed and conducted pinning experiments to validate the theoretical model. JMW designed and conducted the bladder construct experiments with inputs from EEB, XS, ABT, AJH, and MRC. JMW conducted fluorescent staining, and JMW, AL, EEB performed confocal imaging and processing of tissues. ARV performed image analysis with direction from JMW, ABT, AJH, and MRC. JMW, EEB, ARV, and EAS assisted with cell culture. EEB, ARV, and EAS contributed device and hydrogel patterning photos/videos with input from JMW. JMW and AL wrote the manuscript with significant inputs from ABT, AJH, and MRC. All authors have reviewed, edited, and approved of the manuscript.

