## Supporting Information for "Curved by Design: Applying Microfluidic Principles for Nonplanar and Planar Suspended Tissue Patterning to the Development of Bladder Models with Tunable Mechanics"

### Introduction

We first investigate what is the sensitivity of the pinning to a variation of the liquid volume. We consider first the case of a horizontal volume limited by two parallel circular plates with a vertical "feeding" pipette placed in the middle. Finally, the gap (or ring) between the pipette and the solid structure is considered. It is shown that a too small pipette diameter results in depinning probability.

### Part 1: Horizontal device

Let us consider the initial state where the cylinder is filled by liquid, the two open boundaries having "flat" surfaces: the meniscus in the opening with the pipette being flat (zero pressure) and the vertical sides being also in a vertical plane (no deformation), so that the interface is cylindrical (figure 1). This initial state is approximated, because the gravity in the flat cylinder is ignored, i.e. the vertical dimension of the cylinder is supposed small (Bond number $Bo = [\rho g (2\delta)^2 / \gamma]$ <1), and the horizontal curvature of the cylindrical free surface is neglected, i.e. the radius *R* of the cylinder is supposed much larger than the vertical dimension ($R/2\delta \gg 1$).

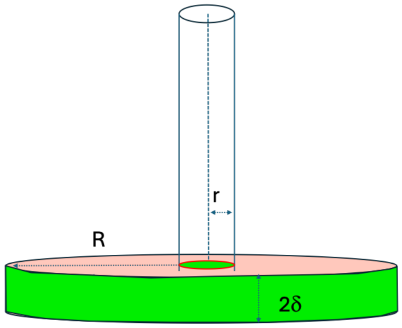


Fig.1: Sketch of the pipette and the disc in the initial state

Now assume that more liquid is added in the pipette. The added volume will contribute to increase the liquid level in the pipette and to trigger bulging of the peripheral free surface (figure 2). Adding sufficient volume of liquid will eventually trigger depinning of the peripheral free surface.

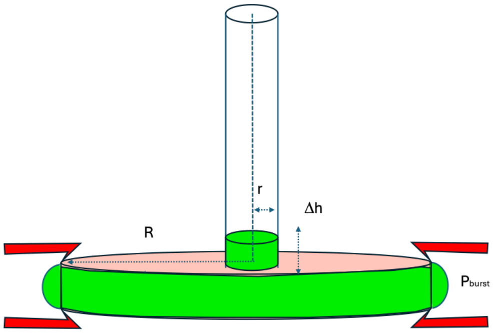


Fig.2: Sketch of the pipette and the disc with the liquid (light green) and the pinning structure (red).

Let us calculate the added volume of liquid in the vertical pipette that triggers depinning. First, we calculate the equilibrium states, when the liquid stays pinned. Consider the two unknowns: $h$ the height of liquid in the pipette, and $R_c$ the curvature radius of the peripheral interface. In the initial state, $h = 0$ and $R_c = \infty$. We have at our disposal two equations linking $h$ and $R_c$: the first one is the pressure equilibrium between the two interfaces. The second one is the splitting of the added liquid volume between the two interfaces.

#### a. Condition #1: pressure equilibrium

The hydrostatic pressure associated to the added liquid in the pipette is

$$P_{hyd} = \rho\ g\ h\ . \tag{1}$$

On the other hand, the pressure of the peripheral free surface (Laplace pressure) is

$$P_{Lap} = \frac{\gamma}{R_c}, \tag{2}$$

where $\gamma$ is the surface tension. The pressure equilibrium relation is

$$P_{hyd} = \rho\ g\ h = P_{Lap} = \frac{\gamma}{R_c}\ . \tag{3}$$

Hence the relation between $h$ and $R_c$

$$h = \frac{\gamma}{\rho\ g\ R_c}\ . \tag{4}$$

#### b. Condition #2: Conservation of volume

The added volume of liquid is the sum of the liquid in the vertical pipette (height *h*) and the bulging volume on the pinning ridge

$$V = V_{pipette} + V_{bulge} = \pi r^2 h + V_{bulge}. \tag{5}$$

Let us calculate the volume $V_{bulge}$. Following the analysis proposed in the book, "The Physics of Microdroplets", let us introduce the total horizontal radius $\tilde{R}$ sketched in figure 3.[1]

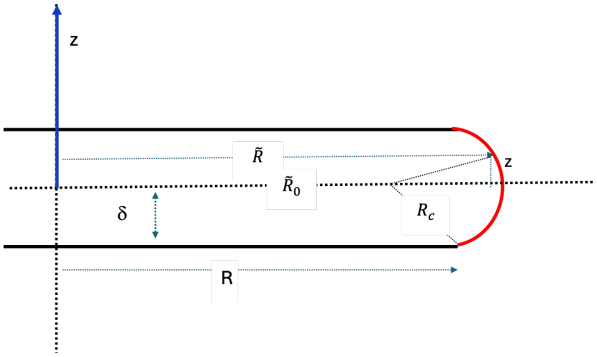


Fig.3. Geometric construction for the calculation of the volume of the bulging liquid

The radius $\tilde{R}$ can be expressed in function of the vertical coordinate *z* by

$$\tilde{R}(z) = \tilde{R}(0) - R_c + \sqrt{R_c^2 - z^2}\,. \tag{6}$$

In (6), $\tilde{R}(0)$ must be specified. To do so, let us express (6) for $z = \delta$ , corresponding to $\tilde{R}(\delta) = R$

$$\tilde{R}(0) = R + R_c - \sqrt{R_c^2 - \delta^2}, \tag{7}$$

Then

$$\tilde{R}(z) = R - \sqrt{R_c^2 - \delta^2} + \sqrt{R_c^2 - z^2}\,. \tag{8}$$

Or, using (4)

$$\tilde{R}(z,h) = R - \sqrt{\left(\frac{\gamma}{\rho g h}\right)^2 - \delta^2} + \sqrt{\left(\frac{\gamma}{\rho g h}\right)^2 - z^2}\,. \tag{9}$$

If we denote $V_{cyl}$ the volume of the cylinder ( $V_{cyl} = \pi R^2 d$), the total volume $V_{bulge} + V_{cyl}$ is given by the integrant

$$V_{bulge} + V_{cyl} = 2\int_0^\delta \pi\,\tilde{R}^2\,dz, \tag{10}$$

so that, after substitution in (5)

$$V = V_{pipette} + V_{bulge} = \pi r^2 h - \pi R^2 d + 2\int_0^\delta \pi\,\tilde{R}^2\,dz, \tag{11}$$

The easiest solution to (10) is a numerical integration. The algorithm is to start from a given height *h,* and integrate (10) by vertical steps $\Delta z$ to obtain the volume *V*.

#### c. Condition #3: Burst pressure and depinning

Adding always more liquid in the pipette results in inflating $V_{bulge}$ unto depinning of the interface when the burst pressure is reached. Let us recall that the burst pressure is determined by[2]

$$P_{burst} = \frac{2\gamma}{d}\sin(\theta + \pi/2\ - \alpha) = \frac{\gamma}{\delta}\cos(\theta\ - \alpha), \tag{12}$$

where $\theta$ is the liquid-solid contact angle and $\alpha$ the pinning structure angle, as shown in figure 4. The pinning is lost when the added liquid volume is such that

$$P_{hyd} = P_{burst}\,, \tag{13}$$

which results in the threshold height in the pipette $h_p$

$$h_p = \frac{\gamma}{\rho g\,\delta}\cos(\theta - \alpha). \tag{14}$$

The corresponding added liquid volume is obtained substituting $h_p$ in (10).

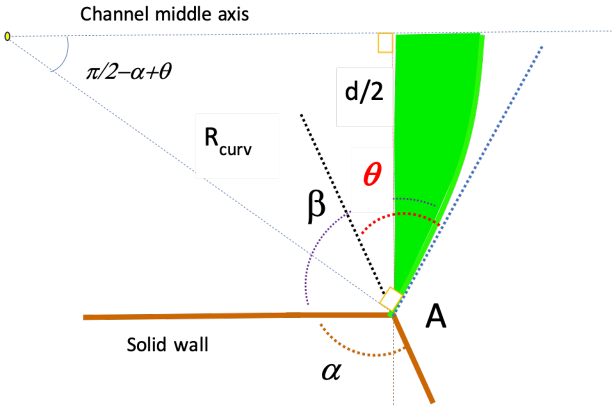


Fig.4. Geometric construction for the location of the meniscus corresponding to the burst pressure where $\gamma$ is the surface tension, $\theta$ the contact angle, $d=2\delta$ the interval between the two plates, and $\tilde{\alpha}$ the structure pinning angle.

### d. Numerical example

Consider the numerical values: $\theta$=35°, $\alpha$ =80°, $\gamma$=30 mN/m, $\rho$=1100 kg/m$^3$, d=2$\delta$=1 mm, $R$=10 mm, $r$=0.8 mm. The results are plotted in figures 5 and 6.

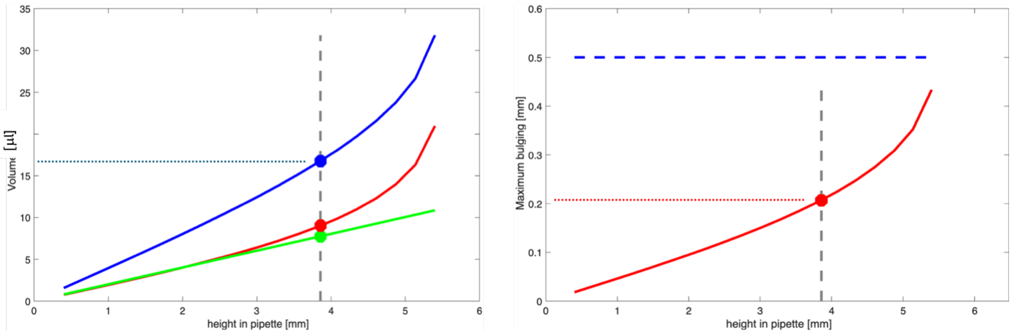


Fig.5. Left: Volumes of liquid as a function of height of liquid in pipette (blue, total volume; red, bulging volume; green, volume in pipette.) Right: Bulging length as a function of height of liquid in pipette. The blue dotted line for the left plot and the red dotted line for the right plot correspond to the depinning limit. The blue horizontal dashed line represents to the maximum bulging possible $d/2$, corresponding to a meniscus in the shape of a semi-circle. The gray vertical dashed lines correspond to where the function intersects with the depinning limit.

The derivative $dV/dh$ can be approximated from the blue curve in figure 5 (left): $dV/dh\sim$ 10 mm$^2$, so that an addition of 1 μl results in an increase of the height $h$ of 100 μm.

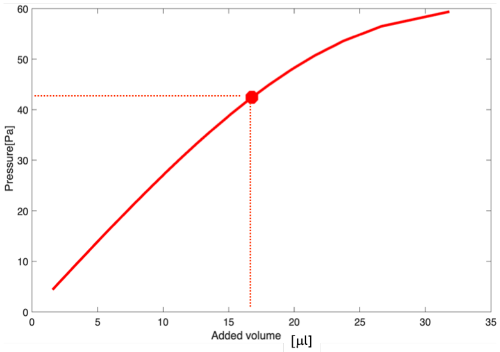


Fig.6. Pressure as a function of the added volume of liquid. The red dotted line corresponds to the depinning limit.

The derivative $dP/dV$ can be approximated from the red curve in figure 6: $dP/dV \sim 1.5$ [Pa/mm$^3$] so that an addition of 1 $\mu$l results in an increase of the pressure of 1.5 Pa.

In figure 7, depinning pressures and liquid heights in pipette are plotted for different contact angles (15 to 40 degrees) and geometric angles (70 to 100 degrees).

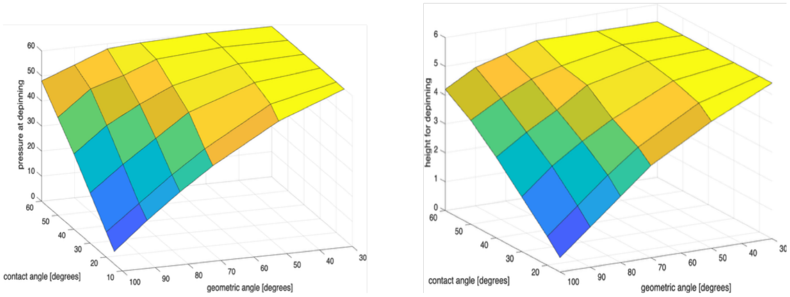


Fig.7. Pinning limits as a function of contact angle and geometric angle, for pressure (left) and liquid height (right).

An analysis of the plots of figure 7 indicates that the pressure at depinning is smaller for larger geometric angles and smaller contact angles. Similarly, the height of liquid in the pipette at depinning is smaller for larger geometric angles and smaller contact angles, as expected.

### e. Algorithm

Let us consider that a volume $V_{tot}$ is placed in the pipette. We use an iterative scheme to find the different volumes: $V_{pip}$, volume in pipette; $V_{bulge}$, bulging out volume of the free interface, and the characteristic lengths: $h_{pip}$, height of liquid in pipette; $R_{bulge}$, radius of curvature of the bulging interface.

The iterative scheme is shown in figure 8.

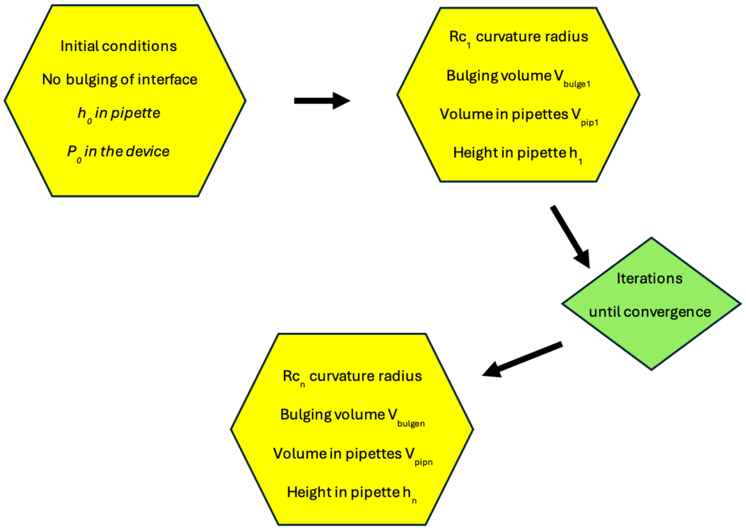


Fig.8. Numerical scheme to obtain equilibrium.

In figure 9, the computed volumes, pressures and characteristic lengths are plotted in function of the radius of the pipette. If the pipette radius decreases (here below 0.8 mm), depinning occurs.

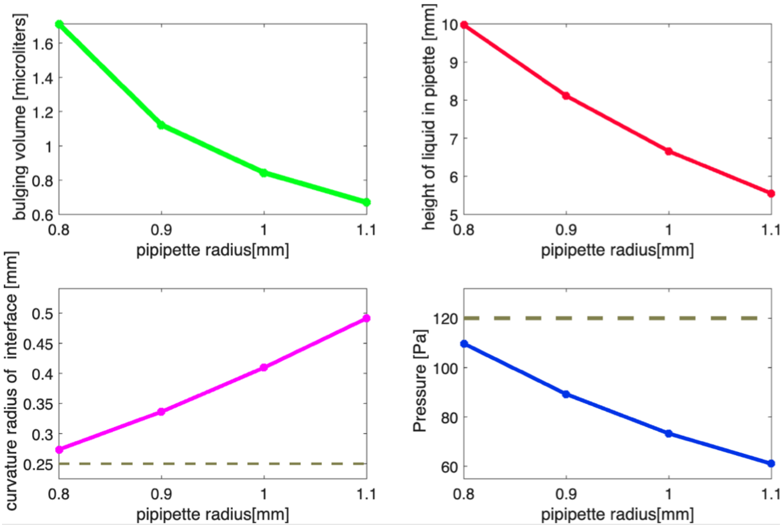


Fig.9. Parametric study as a function of the pipette radius. Top left: bulging out volume; top right: height of liquid in pipette; bottom left: radius of curvature of the free interface; bottom right: pressure of the liquid. The gray horizontal dashed lines correspond to the limit of stability when depinning occurs.

### Part 2: Conical device

Let us now analyze the case of a conical device (referred to as a dome in the main text of the paper), as sketched in figure 10.

#### a. Volumes

Let us calculate the different volumes of interest.

(1) Cone interior

First, calculate the volume in the conical space. The formula for a conical frustrum of height *h* and radii *r* and *R*, is

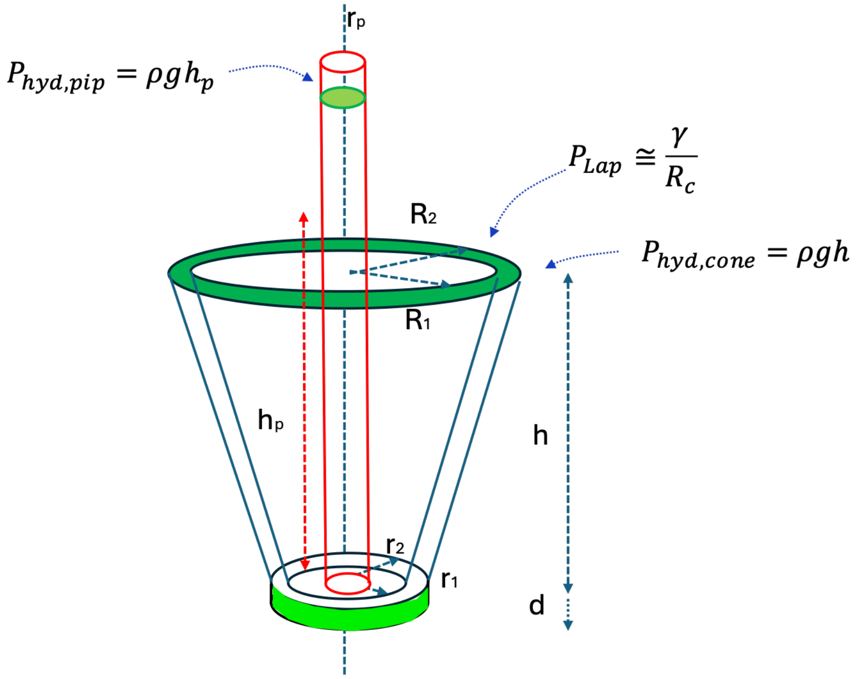


$$v = \frac{1}{3}\pi h\ [(R_2^2 + r_2 R_2 + r_2^2) - (R_1^2 + r_1 R_1 + r_1^2)] + \pi\ r_1^2\ d + \pi\ r_p^2\ h_p$$

Fig.10. Sketch of the device

$$V_{frustrum} = \frac{1}{3}\pi h\ (R^2 + rR + r^2)\ , \tag{15}$$

So that the volume in the conical interspace between exterior radii (index 2) and interior radii (index 1) is

$$V_{cone} = \frac{1}{3}\pi h\ [(R_2^2 + r_2 R_2 + r_2^2) - (R_1^2 + r_1 R_1 + r_1^2)]\ . \tag{16}$$

(2) “Bulging out” volume

The volume “bulging out” at the pinning ridges is a portion of torus, as shown in figure 11. The Pappus of Alexandria’s” theorem is based on the rotation of the cross-surface *S* and yields

$$V_{bulge} = 2\pi \frac{(R_1+R_2)}{2} S, \tag{17}$$

where *S* is a cross surface of the “torus”. This surface S is given by

$$S = \frac{R_c^2}{2} (\alpha - sin\alpha), \tag{18}$$

where the angle $\alpha$ can be expressed in function of $R_c$ as

$$\sin\frac{\alpha}{2} = \frac{(R_2-R_1)}{2\,R_c}. \tag{19}$$

Then (16) becomes

$$V_{bulge} = 2\pi \frac{(R_1+R_2)}{2} \frac{R_c^2}{2} (\alpha - sin\alpha), \tag{20}$$

with $\alpha = 2\,arcsin \frac{(R_2-R_1)}{2\,R_c}$.

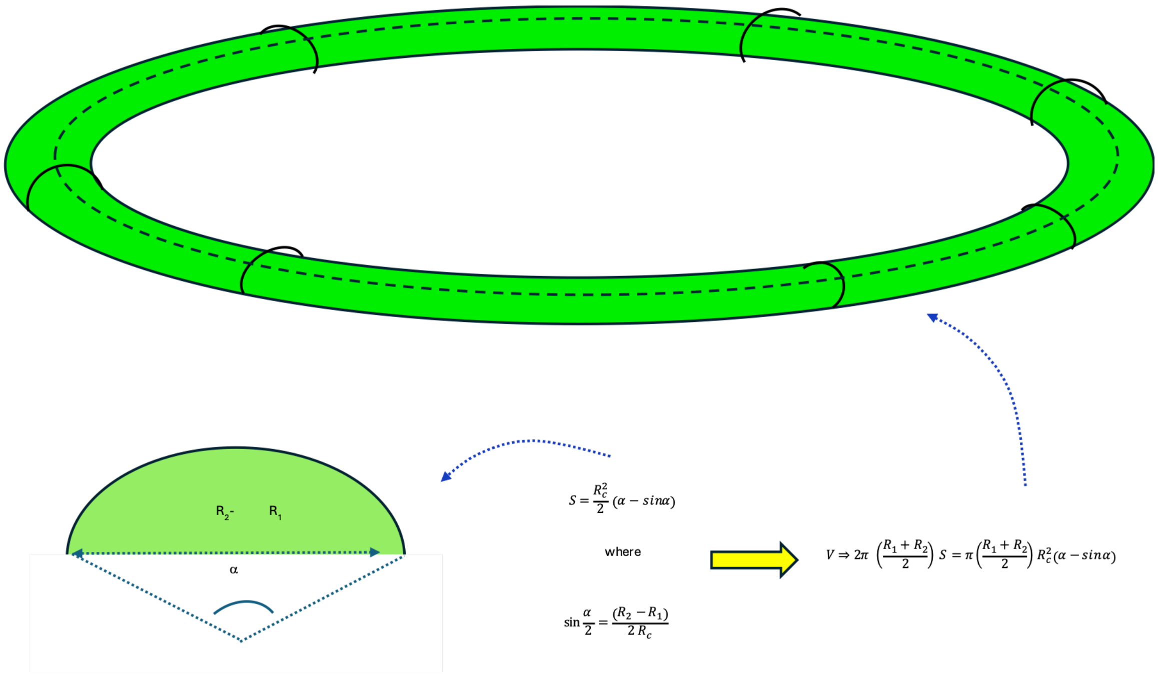


Fig.11: Top: sketch of the bulging surface, with the shape of a portion of torus; bottom: sketch of the cross-surface, *S*.

(3) Basis cylinder

Using the notations of figure 10, the cylinder at the base of the device is

$$V_{cyl} = \pi r_2^2 d. \quad (21)$$

(4) Pipette

$$V_{pip} = \pi r_p^2 h_p. \quad (22)$$

### b. Model

The model is similar to the one for the "flat" case, using the volumes defined above and the hydrostatic pressure $P_{hyd} = P_{hyd,pip} - P_{hyd,cone}$. In table 1, the changes between the two geometries have been listed.

| | notation | "flat cylinder" case | "cone" case |
|---|---|---|---|
| Pressure (hydrostatic) | $P_{hyd}$ | $\rho g h_{pip}$ | $\rho g(h_{pip} - h_{cone})$ |
| Laplace pinning pressure | $P_{Lap}$ | $\gamma/\delta$ | $2\gamma/(R_2 - R_1)$ |
| Volume in the structure | $V_{cyl}$ or $V_{cyl}+V_{cone}$ | $\pi r_{pip}^2 h_{pip} + \pi R^2 d$ | $\pi r_{pip}^2 h_{pip} + \pi r_2^2 d + \frac{1}{3}\pi h\left[(R_2^2 + r_2 R_2 + r_2^2) - (R_1^2 + r_1 R_1 + r_1^2)\right]$ |
| Bulging out volume | $V_{bulge}$ | $2\int_0^\delta \pi \tilde{R}^2\, dz - \pi R^2 d$ | $2\pi \frac{(R_1 + R_2)}{2} \frac{R_c^2}{2} (\alpha - sin\alpha)$ |
| Pipette volume | $V_{pip}$ | $\pi r_p^2 h_{pip}$ | $\pi r_p^2 h_p$ |

Table 1: Differences between the two cases

We use the same algorithm described above using the updated pressures and volumes. The results are plotted in figure 12 for 5 pipette radii: [0.7, 0.8 0.9,1.0, 1.1] mm. The dimensions considered for the calculation are listed in table 2.

| Top interior radius [mm] | Top exterior radius [mm] | Bottom interior radius [mm] | Bottom exterior radius [mm] | Height of pinning ridge [mm] | Semi bottom cylinder height [mm] | exterior cylinder radius [mm] |
|---|---|---|---|---|---|---|
| $R_1$ | $R_2$ | $r_1$ | $r_2$ | h | $\delta$ | $R_{cyl}$ |
| 4.0 | 4.5 | 1.5 | 2.0 | 3.0 | 0.25 | 2.0 |

Table 2: Dimensions of the conical device.

We find qualitatively similar results as that of figure 9 for the geometry of a flat device. If the pipette radius is too small, the hydrostatic pressure becomes larger than the depinning (burst) pressure and the pinning is broken.

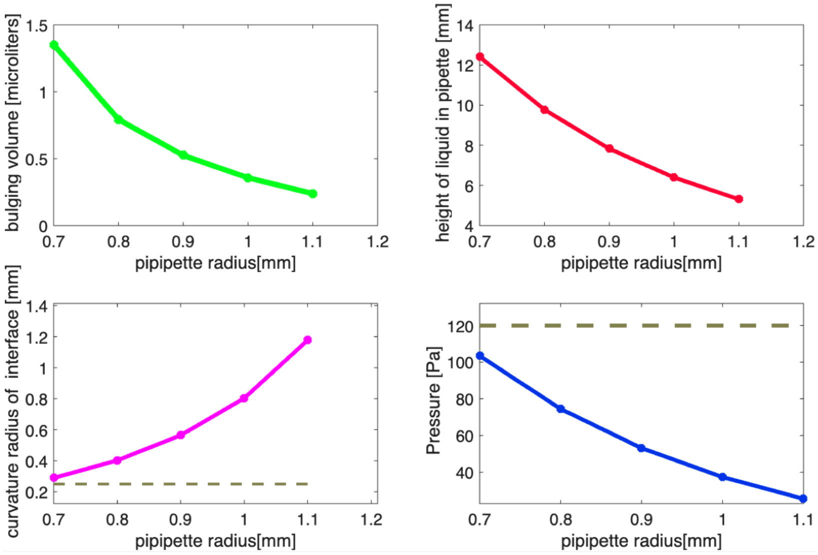


Fig.12: Parametric study as a function of the pipette radius. Top left: bulging out volume; top right: height of liquid in pipette; bottom left: radius of curvature of the free interface; bottom right: pressure of the liquid. The gray horizontal dashed lines correspond to the limit of stability when depinning occurs.

### Part 3: The gap

In the preceding analysis, the gap between pipette and device has been ignored. In this section, the determination of the liquid volume bulging above the gap, as pictured in figure 13, is performed. The surface of the liquid ring is limited by the pinning on the plate edge (blue line in figure 13) and the contact on the vertical pipette (red in the figure). Let us first calculate the height *z* of the liquid along the pipette. The interface curvature radius is given by the hydrostatic pressure in the pipette (neglecting the horizontal curvature):

$$P_{hyd} = \rho g h \cong \frac{\gamma}{R}, \tag{23}$$

Hence

$$R \cong \frac{\gamma}{\rho g h}. \tag{24}$$

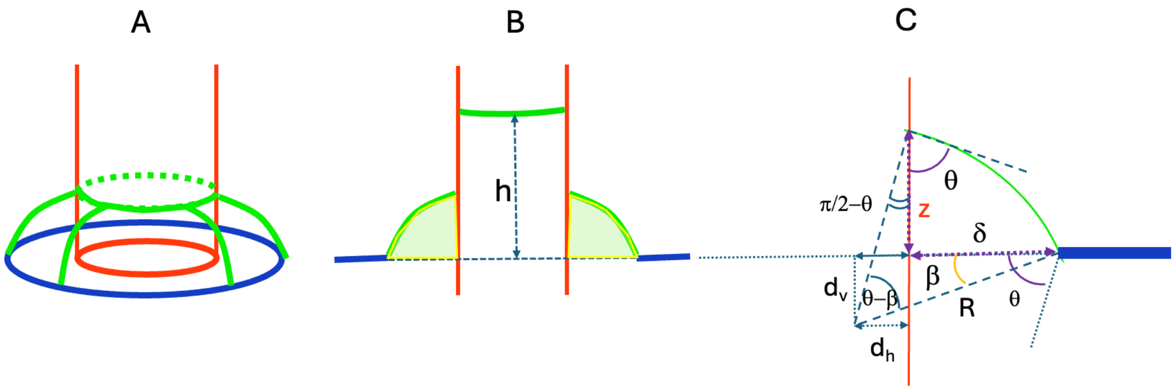


Fig.13. A and B: Sketch of the liquid ring; C: Geometric construction.

Then, referring to the notations of figure 13.C

$$d_h = R \sin\left(\frac{\pi}{2} - \theta\right) = R \cos\theta, \tag{25}$$

and

$$d_v = (d_h + \delta) \tan\beta = (R \cos\theta + \delta) \tan\beta. \tag{26}$$

On the other hand

$$d_v = R \sin\beta. \tag{27}$$

Combining (25) and (26), the angle $\beta$ can be determined

$$\cos\beta = \frac{(R \cos\theta + \delta)}{R}. \tag{28}$$

On the other hand

$$z + d_v = R\cos\left(\frac{\pi}{2} - \theta\right) = R\sin\theta \quad . \tag{29}$$

Finally, the height *z* is given by

$$z = R(\sin\theta - \sin\beta) = R\left[\sin\theta - \sqrt{1 - (\cos\theta + \delta/R)^2}\right]. \tag{30}$$

The surface S in a vertical cross-section is given by the portion of circle plus or minus the surfaces lying outside (shown in figure 14)

$$S = \frac{1}{2}(\theta - \beta)R^2 - \frac{1}{2}d_h\,(z + d_v) + \frac{1}{2}d_h^2\,\tan\beta - \frac{1}{2}\delta^2\,\tan\beta. \tag{31}$$

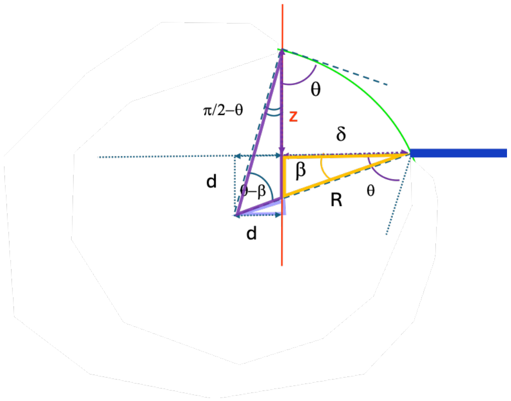


Fig.14. Scheme showing the determination of the liquid surface

Finally, the "Pappus of Alexandria's" theorem based on the rotation of the cross-surface *S* yields the volume of the bulging liquid ring

$$V = 2\pi\, r_{pip} S. \tag{32}$$

The results are shown in figure 15 for a pipette radius $r_{pip}$ of 0.8 mm. The volume of the liquid ring (around 0.1 $mm^3$ or microliters) is relatively small, but not negligible, compared to the volume bulging out from the pinning region (1.0 $mm^3$)

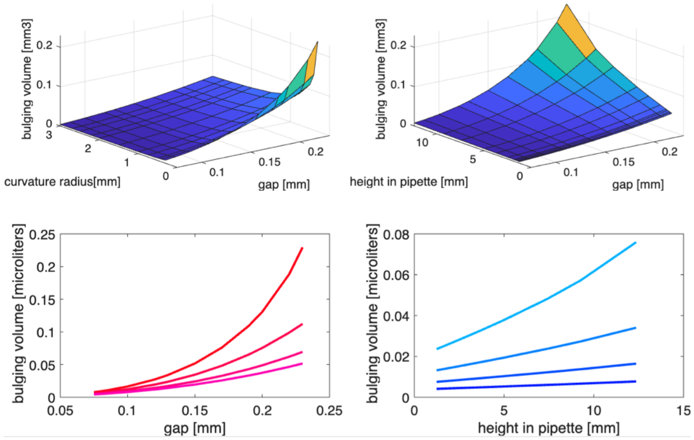


Fig.15. Top left: bulging volume of the ring as a function of curvature radius and gap; top right: bulging volume of the ring as a function of liquid height in pipette and gap; bottom left: bulging volume above ring as a function of gap for different curvature radii; bottom right: bulging volume above ring in function of height in pipette for different gaps.

### Part 4: The whole picture: bulging liquid in the pinning region and gap

The picture of the filling of the device considering the gap (or ring) between the pipette and the solid structure is shown in figure 16.

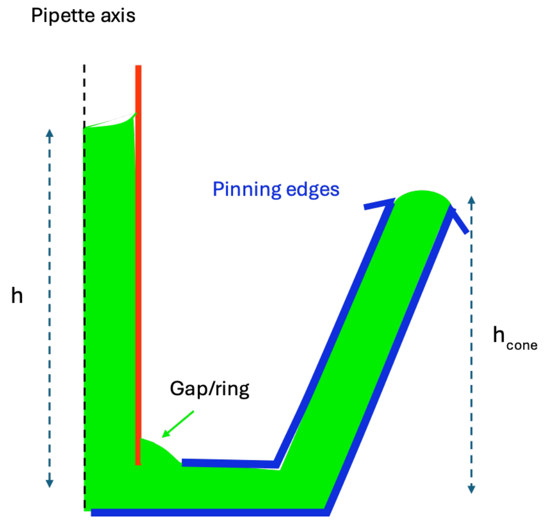


Fig.16. Sketch of the cross section of the 1/2 device with the interface in the pinning region and the bulging in the pipette/structure gap.

The algorithm described in figure 8 has been modified by the presence of the open ring. Figure 17 shows typical results of the model: when the pipette radius is small, the liquid height in the pipette is high and depinning can appear, depending on the device dimensions and liquid contact angles.

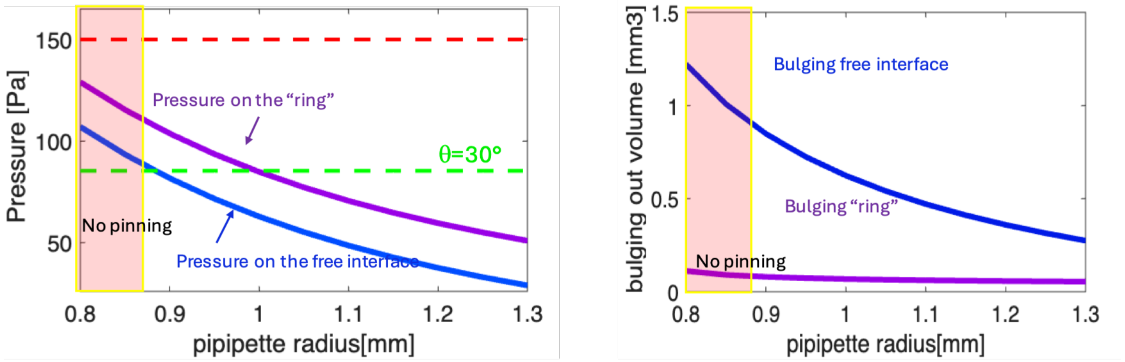


Fig.17. Left: (hydrostatic) pressure on the pinned interface (blue line) and on the open ring surface (purple line); the horizontal red dashed line is the maximum pinning pressure (half spherical interface) and the horizontal green dashed line the actual burst pinning pressure (depending on contact angles); right: volume of bulging liquid as a function of the pipette radius.

### Conclusion

A model for the pinning/depinning of a liquid in a horizontal open cylinder is presented. When the radius of the pipette is increased, the hydrostatic pressure proportionally decreases, the Laplace pressure at the interface at the pinning ridge decreases (at equilibrium it is equal to the hydrostatic pressure), the bulging of the interface decreases, and pinning stability is increased.

The model produces approximate values for the depinning conditions in function of the characteristic dimensions and liquid properties. For a given volume, if the pipette radius is too small, the hydrostatic pressure exerts a burst pressure on the interface and pinning is lost.
